# An image J plugin for the high throughput image analysis of *in vitro* scratch wound healing assays

**DOI:** 10.1101/2020.04.20.050831

**Authors:** Alejandra Suarez-Arnedo, Felipe Torres Figueroa, Camila Clavijo, Pablo Arbeláez, Juan C. Cruz, Carolina Muñoz-Camargo

## Abstract

*In vitro* scratch wound healing assay, a simple and low-cost technique that works along with other image analysis tools, is one of the most widely used 2D methods to determine the cellular migration and proliferation in processes such as regeneration and disease. There are open-source programs such as imageJ to analyze images of *in vitro* scratch wound healing assays, but these tools require manual tuning of various parameters, which is time-consuming and limits image throughput. For that reason, we developed an optimized plugin for imageJ to automatically recognize the wound healing size, correct the average wound width by considering its inclination, and quantify other important parameters such as: area, wound area fraction, average wound width, and width deviation of the wound images obtained from a scratch/ wound healing assay. Our plugin is easy to install and can be used with different operating systems. It can be adapted to analyze both individual images and stacks. Additionally, it allows the analysis of images obtained from bright field, phase contrast, and fluorescence microscopes. In conclusion, this new imageJ plugin is a robust tool to automatically standardize and facilitate quantification of different *in vitro* wound parameters with high accuracy compared with other tools and manual identification.

## Introduction

Cellular behavior regulates wound healing during the phases of proliferation, migration, matrix formation, and contraction. Growth factors and matrix signals that determine the function of cells in regeneration processes orchestrate this behavior (1). Some phases that occur during wound healing have also been observed during cancer invasion (2,3). In order to develop robust therapeutic approaches, it is imperative to study these processes in detail (4).

In this regard, the scratch or migration assay is a widely used tool for *in vitro* studies of the rates of migration (5), angiogenesis (6), movement (7), proliferation (2), and healing in response to different novel drug candidates. Some of these molecules include growth factors, proteins, natural compounds, and small pharmacological principles, among many others (2, 8, 9). This technique is suitable to study the paracrine signals (conditioned media) produced by stem (10) or other types of cells under either mechanical (7) or electrical stimuli (11). Furthermore, it is possible to study the response of free cells and those seeded on scaffolds made of both synthetic polymers and natural extracellular matrices (7, 12–14). In this context, the *in vitro* scratch wound healing assay is useful to evaluate the proliferation and migration process when exposing cells to metabolites present in the conditioned media. This is because this approach provides an environment that mimics that of a wound healing process *in vivo* (15–17). The most remarkable advantages of this assay are the low requirements of specialized equipment or expensive reagents, which makes it adaptable to research groups with limited budgets. Additionally, the produced data is relatively easy to analyze with an ample variety of open access software packages.

This straightforward technique relies on adherent cells that could vary depending on the particular cellular processes to study. Some adherent cell lines used for the assay are endothelial, fibroblast and epithelial (4). The first step is to create manually a “scratch” in a cell monolayer, using pipette tips, razors (17), cell scrapers (18), or any object with a sharp tip. Methods that are more sophisticated include molds or cell inserts (19, 20), electric currents (21), lasers (4) and magnets (20).

The next step is exposure to the treatment and image acquisition at the beginning and at regular intervals during cell migration as the scratch closes. The last step is to monitor the migration path of cells in the leading edge of the scratch by tracking it with the aid of time-lapse microscopy (4) and image analysis software (4, 22).

Some of the most popular software packages for this application include the open source imageJ/Fiji® and licensed ones such as Matlab®. These packages have been previously used to manually count cells (selecting and tallying individual cells) and to assess wound closure (tracing the wound perimeter and calculating the percentage of closure) (23, 24). This approach is however demanding, tedious, and time-consuming. As a result, a number of plugins have been developed for these software platforms to accelerate the analysis process by automatically quantifying cell number and wound area (22, 25). Despite these efforts, existing applications depend on human interaction and require manual measurements, which is problematic for inexperienced users. To overcome these limitations, we designed, implemented, compared, and tested a user-friendly plugin for ImageJ/Fiji®. We tested the plugin in the task of analysing images derived from a scratch assay where Human skin keratinocytes HaCaT were exposed to conditioned media of Human adipose-derived mesenchymal stem cells (hAdMSCs). Our *Wound_healing_size_tool* plugin facilitates a high throughput calculation of a number of parameters from several images including scratch area, wound coverage of total area, scratch width average, and standard deviation of the scratch width.

## Materials and methods

### Reagents

Dulbecco’s Eagle’s medium modified high glucose medium (DMEM) Penicillin/streptomycin (P/S), Fetal bovine serum (FBS), Trypsin-EDTA (1X) were purchased from Biowest (Nuaillé, France) and PBS (1X) pH 7.4 was acquired from Sigma-Aldrich (St. Louis, MO, USA).

### Cell culture

The HaCaT(26) cell line was provided by the Basic Medical Sciences Laboratory of the Faculty of Medicine of the Universidad de los Andes. We grew cells in DMEM supplemented with 10% FBS, 1% P/S at 37 ° C, 5% CO2 and 75% of humidity. We changed the medium three times per week until the cells reached 70% of confluence.

We obtained the hAdMSCs of abdominoplasties realized at Santa Barbara Surgical Center by Dr. Santiago Merchan with previous approval and signing of informed consent and with the approval of the ethics committee of the Vice-presidency of research at the Universidad de Los Andes (Act No 942, 2018). The cells were isolated following the protocol by Linero et al. (27).

### Design of mold for scratching wounds on 2D cultures

We performed mold design using Autodesk Inventor Professional 2020 (Autodesk, Inc., San Rafael, California, USA, www.autodesk.com) to guarantee a completely closed piece with the dimensions of the real object and optimized for laser cutting. The mold was laser cut using a Trotec Speedy 100 CO_2_ (Marchtrenk, Austria). Red lines of 0.1 mm represent the edges cut by the laser cutter, while Blacklines of 0.1 mm delineate regions for engraving. The supplementary information 1 file shows molds for wound scratching for 6, 24, 48 and 96-well plates. We created the schematic representation of the use of the mold for scratching wounds on 2D cultures using (www.biorender.com). Moreover, to compare the scratches made using the molds or using only a 200 μL sterile pipette tip, we evaluated the width homogeneity of the gap with our plugin by the coefficient of variation (CV = SD/mean) and standard deviation with respect to a straight line.

### Wound healing assay

We seeded the HaCaT cells in a 24-well plate at a concentration of 1×10^5^ cells /well and we left them until they reached 80% of confluence (10). We scratched cell cultures with a 200 μL sterile pipette tip (7,22) using the designed mold (see above for details) (Supplementary information 1). We then washed away detached cells with PBS (1X). Then, we added 1 mL of hAdMSCs conditioned medium, which was obtained from cells in the 4-6th passage according to the protocol established by Qazi et al.(7). We made horizontal reference lines on the bottom of the plate with an ultrafine tip marker to have a grid for alignment to obtain the same field for each image acquisition run. Once we made the reference lines (approximately 3000 µm of distance), the plate was placed under a phase-contrast microscope using as guide the reference marks. We analyzed selected regions of interest using a Zeiss inverted microscope on a 10X objective with NA/0.25 every 2 hours for 28 hours. We determined the scratch area, wound coverage of total area, and average and standard deviation of the scratch width with the aid of our plugin. We calculated the rate of cell migration (R_M_) and the percentage of wound closure according to (Eq 1) and (Eq 2), respectively (28):

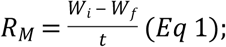

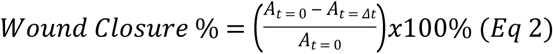

Where W_i_ is the average of the initial wound width, W_f_ is the average of the final wound width both in µm and t is the time span of the assay in hours. Additionally, *A*_*t*=0_ is the initial wound area, *A*_*t*=*Δt*_ is the wound area after n hours of the initial scratch, both in µm^2^.

### Wound healing size plugin for ImageJ®

Inspired by the Wound Healing Tool by Montpellier Resources Imagerie, we designed *Wound Healing Size Tool*, an ImageJ/Fiji® plugin that allows the quantification of wound area, wound coverage of total area, average wound width and width standard deviation in images obtained from a wound-healing assay.

To discriminate adequately between cell monolayer and open wound area, we developed a classic computer vision segmentation algorithm focused on assessing neighboring pixel intensity variance (i.e., the morphology and visual characteristics). With careful inspection of the gray-scaled images obtained from the wound healing assay, we noticed that the pixels in open wound regions had very similar intensity values, while cell monolayer regions had higher variation due to cell presence. Accordingly, analyzing the variance within different neighborhoods in the image could be helpful for discriminating between cell monolayer and open wound regions. To do this, we initially enhanced the contrast of the image to increase the variance within the cell monolayer and facilitate its posterior binarization. Subsequently, we applied a variance filter (by making use of Image J built-in macros *Enhance Contrast* and *Variance filter*), which calculates the intensity variance in the immediate neighborhood of each pixel, and labels it with its corresponding neighborhood variance. This creates a new image with high intensity pixels within the cell monolayer and low intensity pixels within the open wound area. Therefore, we can apply a simple threshold to the processed image in order to obtain binary segments with a user-defined input. However, since the open wound area usually contains single cells or cell islets, the obtained binary image will not include these regions as part of the wound. To remove these structures, we perform a morphological reconstruction by erosion, also known as *hole filling*, on the area binary labeled as open wound. This operator detects all connected components enclosed by the wound area and it includes them as part of the wound. Moreover, since there can be small regions within the cell monolayer with low variance, this algorithm can yield more than one region labeled as open wound. To eliminate the regions falsely detected as wounds, we select the largest connected component within the detected open wound regions as the true wound area. Figure 1 presents an overview of our image-processing algorithm.

**Figure 1.**
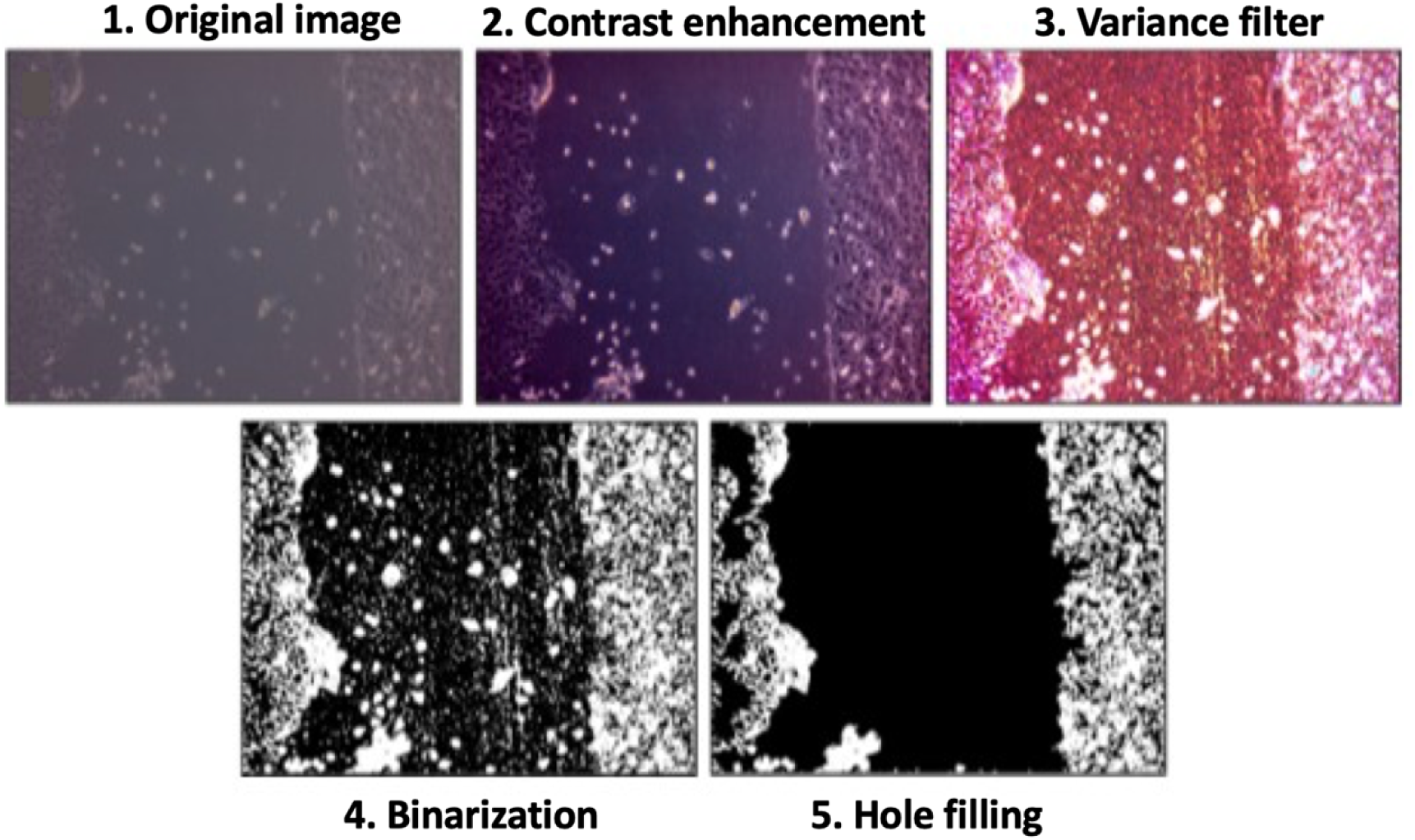
Overall process of the image segmentation algorithm. We show images from each of the main steps to illustrate how the wound area is separated from the cell monolayer area.

After segmenting the open wound, its area and the coverage with respect to the whole image are easily determined with the region’s metadata using the ROI manager. However, for wound width analysis, we quantify the distance between the lateral wound edges at every y-coordinate. Accordingly, we identify edge pixels within the segmented region and we pair them according to their y-coordinate. We calculate the Euclidean distance between paired pixels, and we determine their average to describe wound width. Moreover, we also consider the standard deviation of the calculated widths to assess the homogeneity of the scratch and subsequently its closure. In addition, the algorithm also considers cases in which the wound is not vertical but made at a specific angle. In these cases, it determines the inclination angle and adjusts each calculated width with trigonometric relations.

This plugin is useful for analyzing wound images since it relies on user-defined input values for the neighborhood radius of the variance filter, the threshold value for binarization, and the saturation percentage in the contrast enhancement, which can vary depending on the analyzed image. The variance window radius represents the radius of the variance filter, which we establish to determine the empty or the occupied zones. The radius must be big enough, so that the noise variance has no impact on tissue variance. The percentage of saturated pixels allows enhancing the contrast of the image by determining the number of pixels that could saturate in the image. By increasing this value, we expect an increase in the contrast. This value should always be greater than zero. Finally, we convert the image resulting from the variance filter to a mask by applying the given threshold.

The plugin features an interface window with two options to adjust the values of all parameters (Figure 2A). The first one applies when the user has multiple images with the same calibration scale. The second one considers that if the scratch is diagonal, with fixed width measurement according to the inclination angle, derived by fitting the selected ROI to an ellipse. The plugin presents all results for each image, area of the wound, wound coverage of total area, and average of the width and its standard deviation in a table with the set scale (Figure 2B and 2C). This is for both Individual image analysis and Stack analysis. The plugin and detailed information and user manual can be found in the Supplementary information 2 file and in https://github.com/AlejandraArnedo/Wound-healing-size-tool/wiki as an open source wiki. To validate the quality and efficiency of Wound_healing_size_tool, we compared it with other ImageJ plugins, including MRI_Wound_Healing_Tool (http://dev.mri.cnrs.fr/projects/imagejmacros/wiki/Wound_Healing_Tool) (MRI), the ScratchAssayAnalyzer tool from A microscope image analysis toolbox (MiToBo) (https://mitobo.informatik.uni-halle.de/index.php/Applications/ScratchAssayAnalyzer) (29), as well as manually drawn edges of the scratch (Manual) in 30 different images. We quantified the differences between existing methods regarding our plugin using Eq. 3:

**Figure 2.**
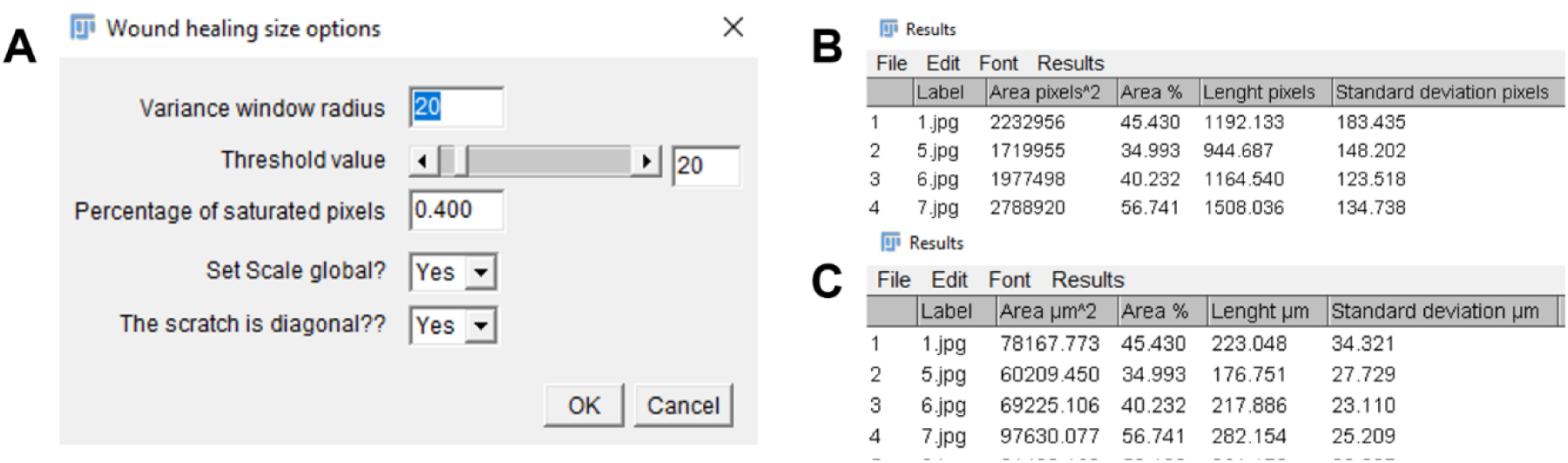
Wound healing size tool. **A. Interface** window to adjust parameters **B.** Snapshot of the results in table format in pixels show area of the wound, wound coverage of total area, and average of the width and its standard deviation**. C.** Snapshot of the results in table format in µm show area of the wound, wound coverage of total area, and average of the width and its standard deviation.

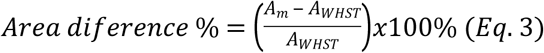

Where A_m_ is the area in pixels calculate using the other ImageJ plugins (MRI, MiToBo) and the manual method, Where A_WHST_ is the area in pixels calculate using our plugin Wound_healing_size_tool (WHST),

### Statistical analysis

We performed Paired t-tests to compare the difference between the area and the average length obtained with the Wound_healing_size_tool with respect to the manual measurement and the other available macros for ImageJ/Fiji®, considering as significant p-values of less than 0.05. We analyzed and plotted The results using GraphPad Prism software 7 (GraphPad Software, San Diego, California, USA, www.graphpad.com).

## Results

### Validation of Wound formation mold

The manually raised hand induced scratch is highly variable with respect to the width, and this variability affects the cell migration rate and the reproducibility of the assay (17, 20, 28). To address this issue, we designed an acrylic (polymethyl methacrylate) mold that fits 24, 48, and 96 well plates (supplementary information 1) (Figure 3A). The results of the width homogeneity of the scratchFigure 3B), using the mold, show a reduction in variability of approximately 50% when compared with manually made scratches using a pipette tip. Although both methods require the use of a pipette tip, the mold gives the advantage of limited tip movement and fixed inclination for more uniform scratch edges. The variation was only of about 5% (Figure 3C).

**Figure 3.**
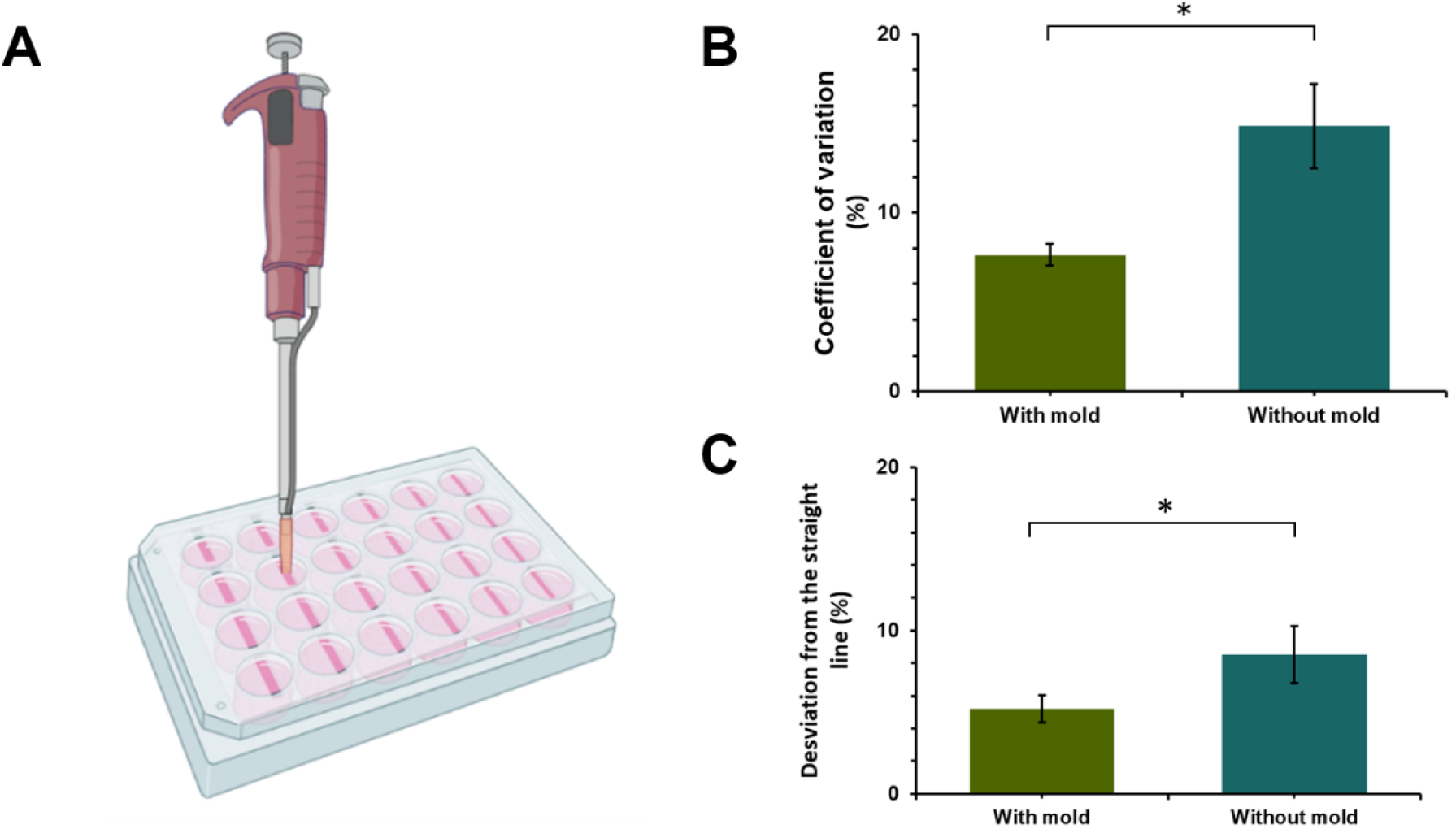
Scratch validation using mold. **A** Schematic of the use of mold for scratching wounds on 2D cultures**. B.** Percentage of coefficient of variation between scratch made with pipette tip of 200 µL with or without using the mold [n=4 images per method]. **C.** Percentage of deviation from the straight line of each scratch made with pipette tip of 200 µL with or without using the mold [n=4 images per method] p_value<0.05*.

### Wound healing size plugin evaluation

Before running the analysis in all the stack of collected images, we selected three images randomly at different time points to set the right parameters for wound identification. We evaluated the performance of our plugin by analyzing 60 wound images. Then, we completed the stack analysis and, with the resulting data, we obtained other parameters such as the closure area percentages, and the rate of cell migration at each time point (Figure 4).

**Figure 4.**
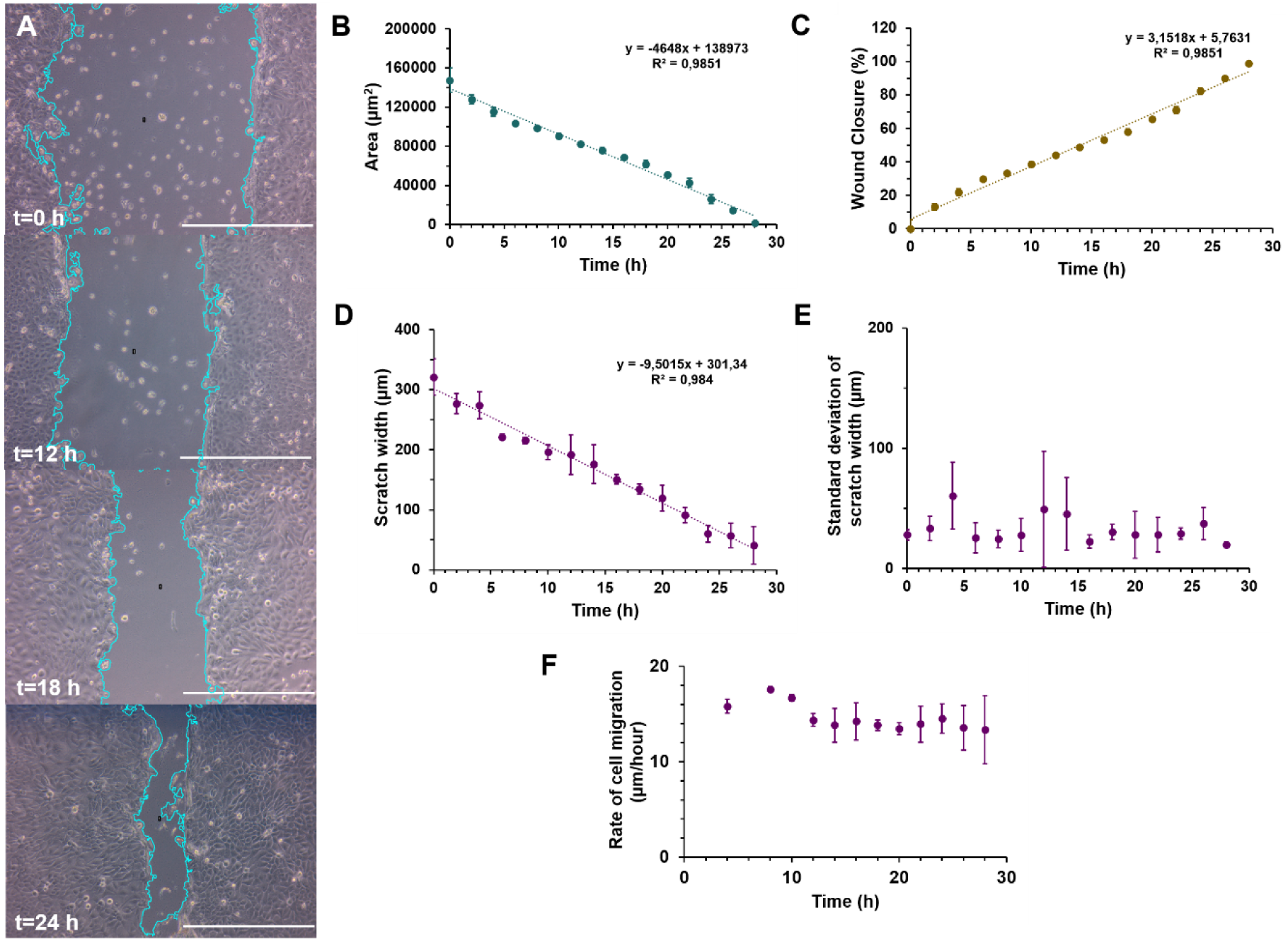
Wound healing size tool test. **A.** Wound healing assay in HaCaT exposed to hAdMSCs conditioned medium at 0, 12 18 and 24 hours. Scale bar=250 µm. **B.** Scratch area in µm^2^ **C.** Percentage of closure area **D.** Scratch width in µm**. E.** Standard deviation of the scratch width in µm**. F.** Rate of cell migration in µm/hour. All the measurements and parameters were taken for 28 hours [n=4 replicas per time]. Some error bars are shorter than the size of the symbols.

As shown in Figure 4, conditioned medium from hAdMSCs promoted healing in an in vitro epidermal model. According to Figure 4B and 4C, the wound closure proceeds at a constant rate, without significant changes over the period evaluated (R^2^=0.9851) reaching almost 95% of closure after 28 hours. However, Figures 4D and 4F demonstrate how the rate of cell migration is higher during the first eight hours of the assay with values that approached 18 µm/hour. In addition, we can observe that, after 12 hours, the rate of cell migration approaches an average value of 13.5 µm /hour until the end of the experiment. Finally, Figure 4E shows the standard deviation of the scratch width, which is an indicative of the homogeneity of the scratch width at each time and it provides insights into how the cells migrate. At the initial time, the standard deviation is lower when compared with later times. This is likely due to the impact of the scratching method on the standard deviation at the outset of the experiment. We observe a higher standard deviation after 12 hours, which is the approximate time when a major change in the cellular behavior occurs and, consequently, the rate of cell migration decreases.

## Discussion

### Validation of Wound formation mold

Two of the principal attributes to be found in a standardized assay are reproducibility and repeatability. Reproducibility in the scratches is imperative, as it has been reported to affect significantly cell migration rates (17, 28). Nonetheless, the implementation of the wound healing assay is generally challenging due to the impact of multiple variables such as the tool used to make the scratch, the pressure, the velocity of the scratching, and the experience of the researcher (19,20,30). For these reasons, we designed a polymethyl methacrylate mold for a 24-well plate. This mold is reusable and can be sterilized using different strategies such as supercritical carbon dioxide, hydrogen peroxide gas plasma, ethylene oxide, and even γ-irradiation (31, 32). Furthermore, the mold produced more reproducible and homogeneous scratches in contrast with gaps made using only a pipette tip. This is possible due to the reduced movement of the pipette tip enabled by the mold, which prevented inclination and helped to control the velocity and the pressure applied to make the scratch. Additionally, our plugin allowed us to compare easily the two methods evaluated to make the wounds.

### Wound healing size plugin evaluation

As shown in Figure 4, the exposure of hAdMSCs to the conditioned medium promotes the migration and wound healing of keratinocytes. These results are consistent with previous studies that have established the potential of mesenchymal stem cells secretome in cell regeneration and migration (7, 10, 27, 33, 34). This has been attributed to the components of the medium, which include multiple growth factors such as Transforming growth factor-beta one (TGF-β1), Epidermal growth factor (EGF), Fibroblast growth factor (FGF), Hepatocyte growth factor (HGF), Interleukin 6 and 8 (IL-6 and IL8), as well as components of the extracellular matrix such as type I collagen and fibronectin (10). All these biomolecules activate signaling cascades that enhance the migratory and proliferative potential of various cell types (7, 10, 35). The wound closure values after 12 hours (Figure 4C) were higher compared with the results reported by Walter et al., (higher than 20%) but consistent than those found by them after 28 hours (closer to 95%) (10). This result is most likely due to the high rate of cellular migration (aprox. 18 µm/hour) during the first 12 hours of the experiment. Nonetheless, after this time, we reduced the migration rate to 13.5 µm/hour (Figure 4F). We estimated these migration times with the data analysis provided by our plugin. Besides analyzing specific points in time, the plugin can be useful for time-lapse analysis and even the variability and heterogeneity of the cellular migration. Importantly, the time for analysis for 60 images was only 5 to 8 min, which represents less than 10% of the time it takes to do it manually.

### Comparison with other imageJ plugins and the manual analysis

To validate the accuracy of our plugin to locate the scratch in each image, 30 images were analyzed and compared with the MRI wound healing tool, the ScratchAssayAnalyzer in MiToBo and with manually drawing the scratch. The area obtained with our plugin led to no significant differences with respect to the other methods (Figure 5). Figure 5A shows how all evaluated methods distinguish gaps with regular shape similarly. Moreover, in some regions, the detected edges of the scratch area overlapped between methods. In contrast, Figure 5B shows that for gaps with an irregular shape, even though we used the same parameters of the MRI method, our plugin determined an area that is closer to that of MiToBo. This can be due to the image enhancement used as pre-processing. Nonetheless, changing the input parameters in our plugin alters the detected area. Accordingly, it is advisable to test two or three images with different combinations of the parameters before starting the analysis in order to localize an appropriate area of analysis.

**Figure 5.**
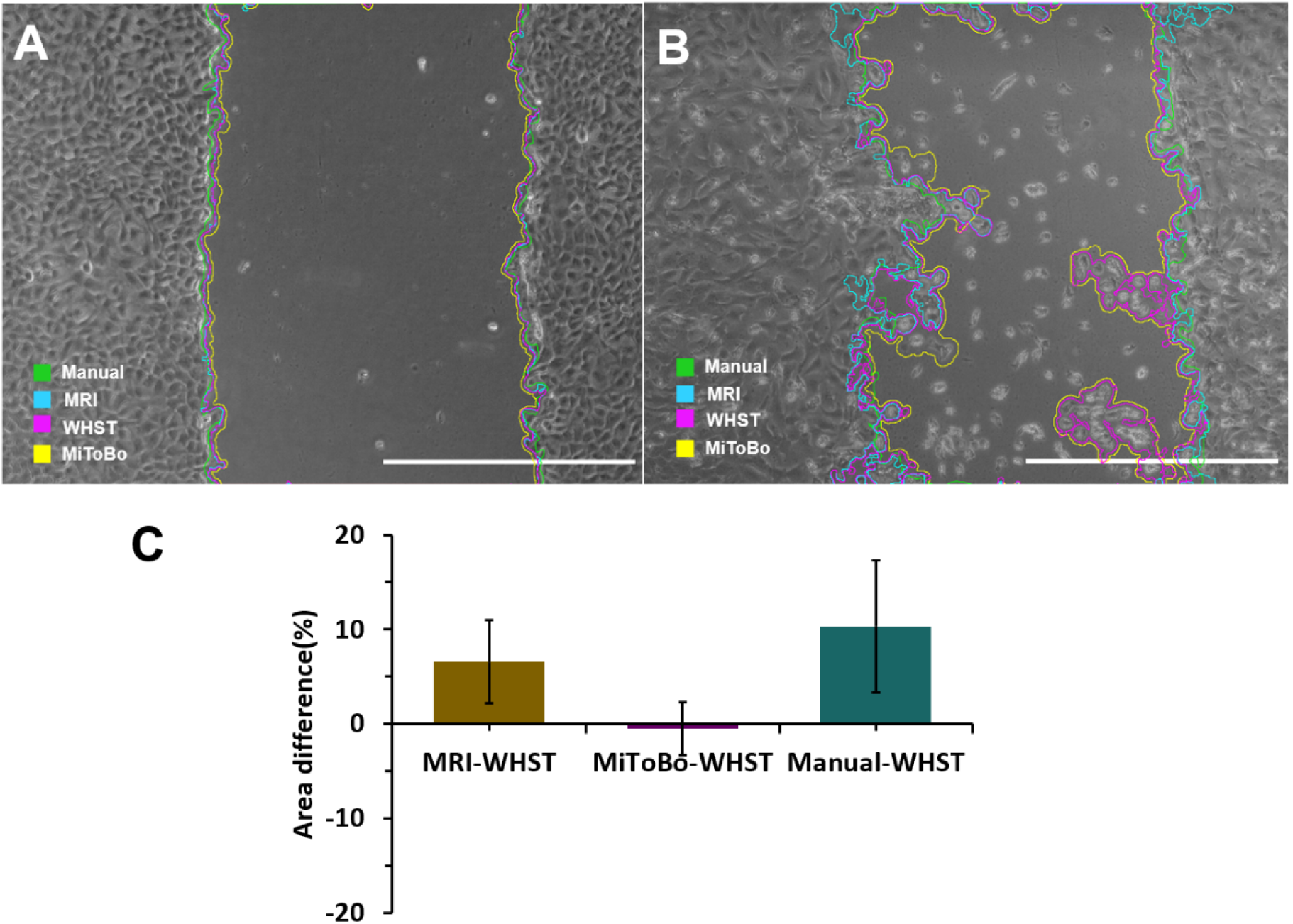
Differences in the area measured with the wound healing size tool (WHST) regarding the MRI wound healing tool (MRI), the ScratchAssayAnalyzer in MiToBo - A microscope image analysis toolbox (MiToBo) and manual drawing (Manual). **A.** Scratch with low variation of calculated area between the different methods. Scale bar=1000 pixels **B.** Scratch with high variation of calculated area between the different methods **C.** Quantification of the difference in measurements between methods with our plugin [n=30 images].

According to Figure 5C, our plugin, which uses a variance-based method, and after enhancing contrast, estimated an area similar to that obtained with the ScratchAssayAnalyzer in MiToBo (only -0.51%±2.8 of difference). This approach is considered highly reliable as it relies on a user-trainable and sophisticated support vector machine (SVM) algorithm (29). With respect to the MRI wound healing tool (which is also based on a variance-based method (19)), our plugin also underestimated the area by 6.5%± 4.4. The average runtime per image is another plugin feature we evaluated. In this regard, our plugin is comparable to the MRI wound healing tool as they both take between 6 to 13 seconds to perform the image analysis. In contrast, since the average execution time of the ScratchAssayAnalyzer in MiToBo is about 2 minutes our plugin is 94% faster.

None of the available plugins for ImageJ calculates the width of the wound, and therefore the comparison was only made with respect to the manual method. According to Figure 6, there is no significant difference between the average width calculated with the plugin or calculated manually. Nonetheless, there is a significant difference (of about 36%) in the standard deviation between the two methods. This is most likely because, as described by Liang et al. (17), at least 100 measurements of width are required to have a proper standard deviation estimate.

**Figure 6.**
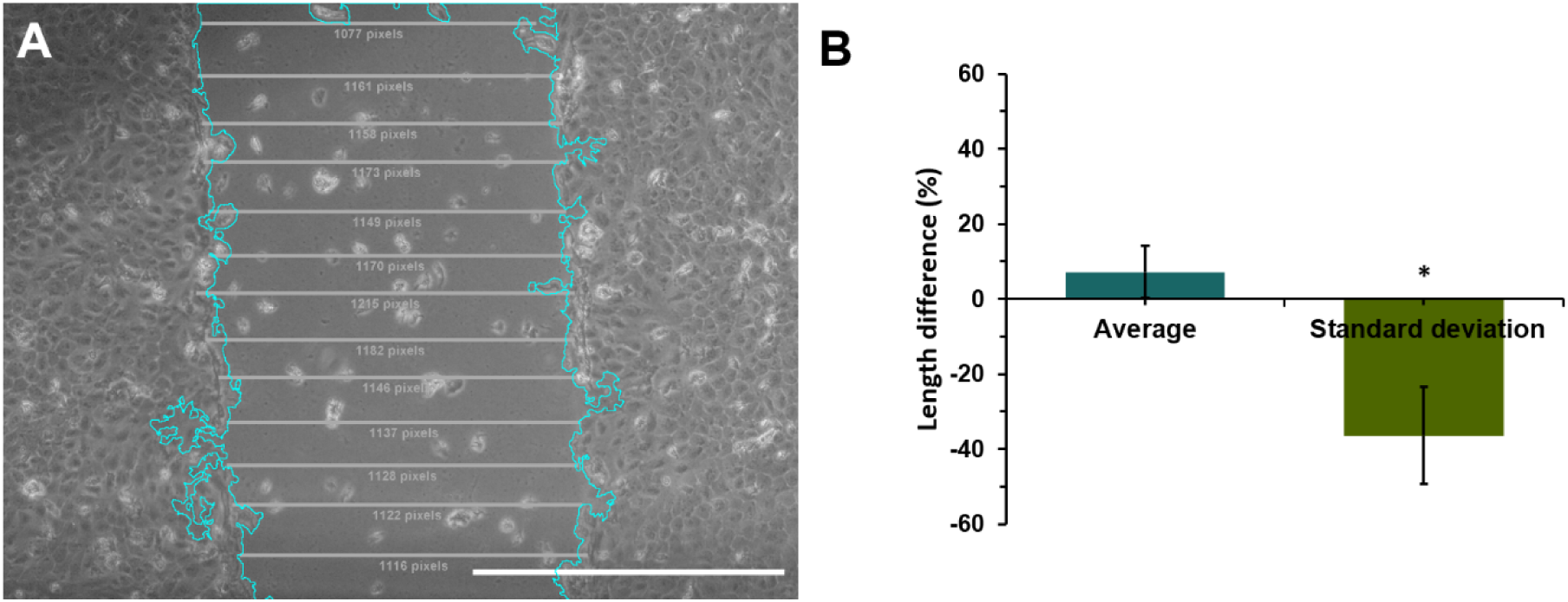
Differences in the width measured with the wound healing size tool (WHST) regarding the manual drawing (Manual). **A** Scratch identifies with our plugin. Wound with manual measurement of width. Manual data (average =1149 pixels, standard deviation =22.82 pixels) WHST data (average =1165 pixels, standard deviation 102.35 pixels). Scale bar=1000 pixels **B.** Quantification of the difference in measurements between wound healing size tool (WHST) and scratching manually [n=30 images, 10 lines per area in the manual method] *p_value<0.01.

The measures of the wound’s width is likely to be impacted by its inclination. For this reason, we included a function in the plugin that allows correction with respect to an inclination angle. An analysis of the same wound but with different inclination showed an average width difference of only 1.35 + -0.59% with respect to the original measurement, which was perpendicular to the x-plane of the image (Figure S1). This proves the effectiveness of our plugin to calculate the wound’s width for different imaging conditions. Finally, we evaluated the accuracy of the plugin to detect the wound area in images collected in bright field, phase contrast, and fluorescence microscopes. The image analysis allowed us to validate that the plugin can be useful in images collected in a wide variety of instruments without introducing significant changes in estimated parameters (Table S1 and Figure S2), in contrast to some macros that only work to identify the scratch from images collected with a limited number of instruments (25).

## Conclusion

The wound healing size tool provides an automatic method of processing images obtained from a scratch/ wound healing assay. This plugin facilitates the quantification of wound characteristics and circumvents their manual acquisition in many images obtained during each experiment. The wound healing size tool has the following advantages:

- It quantifies the area, wound coverage of total area, average wound width and width standard deviation.
- It is easy to use and can be adapted to analyze both individual images and stacks.
- It saves analysis time
- It can be used with different operating systems and it is easy to install.
- It corrects the average wound width by considering its inclination
- It allows the analysis of bright field, phase contrast, and fluorescence images.

## Acknowledgments

This work was financially supported by the Department of Biomedical Engineering at Universidad de los Andes and the Fondo de Apoyo a Profesores Asistentes of Carolina Muñoz. We would like to acknowledge Natalia Bolaños and Dr. John Mario González for providing the HaCaT cells for the study. Additionally, special gratitude to Laura Bocanegra for her collaboration with image acquisition.

## Supporting information captions

- **Supplementary information 1.** Blueprints Wound formation molds for 6, 24, 48 and 96-well plates
- **Supplementary information 2.** Manual Wound_healing_size_tool (Individual image analysis) and Manual Wound_healing_size_tool (Stack analysis).
- **Figure S1.** Fixing the width of the wound with respect to its inclination. A. Wounds with different angles of inclination. Scale bar=200 pixels. B. Width measurements with (blue) and without correction (white).
- **Table S1.** Parameters set per image.
- **Figure S2**. ROI selection in images analyzed with different set parameters (Table 1).

